# Source Localization Using Recursively Applied and Projected MUSIC with Flexible Extent Estimation

**DOI:** 10.1101/2023.01.20.524892

**Authors:** Lukas Hecker, Ludger Tebartz van Elst, Jürgen Kornmeier

## Abstract

Magneto- and electroencephalography (M/EEG) are widespread techniques to measure neural activity *in-vivo* at a high temporal resolution but low spatial resolution. Locating the neural sources underlying the M/EEG poses an inverse problem, which is ill-posed. We developed a new method based on Recursive Application of Multiple Signal Classification (MUSIC). Our proposed method is able to recover not only the locations but, in contrast to other inverse solutions, also the extent of active brain regions flexibly (FLEX-MUSIC). This is achieved by allowing it to search not only for single dipoles but also dipole clusters of increasing extent to find the best fit during each recursion. FLEX-MUSIC achieved the highest accuracy for both single dipole and extended sources compared to all other methods tested. Remarkably, FLEX-MUSIC was capable to accurately estimate the level of sparsity in the source space (*r* = 0.82), whereas all other approaches tested failed to do so (*r* ≤ 0.18). The average computation time of FLEX-MUSIC was considerably lower compared to a popular Bayesian approach and comparable to that of another recursive MUSIC approach and eLORETA. FLEX-MUSIC produces only few errors and was capable to reliably estimate the extent of sources. The accuracy and low computation time of FLEX-MUSIC renders it an improved technique to solve M/EEG inverse problems both in neuroscience research and potentially in pre-surgery diagnostic in epilepsy.

## 2 Introduction

In this paper, we present a novel approach for solving the inverse problem of magneto and electroencephalography (M/EEG) using truncated recursively applied multi-signal classification for sources with variable coherent (FLEX-MUSIC). The EEG and MEG inverse problem is a fundamental challenge in the field of neuroscience, as it involves inferring the underlying neural activity that generates a given set of EEG or MEG measurements (Awan, Saleem, & Kiran, 2019; He, Sohrabpour, Brown, & Liu, 2018; Michel, n.d.; Michel & Brunet, 2019; Nunez & Srinivasan, 2006). The problem is, that many different configurations of brain activity can cause the same signal measured on the scalp. Traditional methods for solving this inverse problem rely on mathematical assumptions that are often not aligned with biophysical models of the brain. Others rely on statistical techniques or optimization algorithms, which can be computationally expensive and may not scale well to large datasets. Despite the underdetermined nature of the M/EEG inverse problem, inverse solutions help researchers and clinicians alike. M/EEG inverse solutions help researchers gain insights into the spatio-temporal workings of the brain during, e.g., perceptual and/ or cognitive processing (Feige et al., 2005; Kornmeier, Friedel, Hecker, Schmidt, & Wittmann, 2019; Luck, 2014). It allows for the identification of functional brain networks and is also used in neurofeedback applications where participants learn to control brain activity in different frequency bands within a specified location (van Lutterveld et al., 2017). Clinicians utilize M/EEG inverse solutions to categorize brain abnormalities or to guide brain surgery, e.g., in drug-resistant epileptic patients (Aydin et al., 2015; Ebersole, 1994; Lantz, Holub, Ryding, & Rosén, 1996; Sharma et al., 2018; Willemse, Hillebrand, Ronner, Peter Vandertop, & Stam, 2016).

There are several existing approaches for solving the M/EEG inverse problem (henceforth referred to as *solvers*), each with its own strengths and limitations. A popular class of methods are the minimum norm estimates (MNE, Hamalainen, 1984; Hämäläinen & Ilmoniemi, 1994; Pascual-Marqui, 1999) that aim to find a solution that explains the observed EEG data with minimal energy of the sources. While these approaches typically incorporate the L2-norm of the source, L1-type solvers have been proposed Beck and Teboulle (2009) and termed minimum current estimates (MCE) in the domain of M/EEG inverse problems. These L1-type solvers find sources that are sparse in nature since the imposed L1-penalty ultimately sets most dipole values close to zero.

Low-resolution tomography (LORETA) and its iterations, standardized and exact LORETA, fall within this class of solvers (Pascual-Marqui, 1999, 2002, 2007). While these approaches are fast to compute and easy to interpret, these approaches often produce blurred solutions, despite their ability to correctly localize source maxima.

Beamformers are a class of spatial filtering techniques that are commonly used to solve the M/EEG inverse problem. These methods involve constructing a set of spatial filters that are applied to the measured signals to estimate the neural activity at each location in the brain. Beamformers can be effective in certain scenarios but struggle to correctly localize multiple correlated sources or spatially coherent source activity. A commonly used proponent is the linearly constrained minimum variance (LCMV) Beamformers (Grech et al., 2008; Van Veen, van Drongelen, Yuchtman, & Suzuki, 1997). A novelty in the field of Beamformers are the multiple constrained minimum variance (MCMV) Beamformers, which alleviate the problem of correlated sources (Nunes et al., 2020).

Bayesian methods provide a framework for incorporating prior knowledge and uncertainty into the inverse solving process (Friston et al., 2008; Grech et al., 2008; D. Wipf & Nagarajan, 2009; D. P. Wipf, Owen, Attias, Sekihara, & Nagarajan, 2010). These approaches involve constructing a probabilistic model of the forward and inverse problems, and then apply Bayesian inference to estimate the posterior distribution of the neural activity given the measured M/EEG signals. Bayesian approaches, like sparse Bayesian learning (SBL, Friston et al., 2008; D. Wipf & Nagarajan, 2009), are capable to accurately localize sources under different sparsity assumptions. However, Bayesian optimization can often be computationally intensive. This problem is amplified when many dipoles are present in the source space, exacerbating the computational load.

A novel class of inverse solvers arose in the past decade that utilize the recent advances in machine learning, predominantly artificial neural networks (ANNs) to solve M/EEG inverse problems. These approaches require training an ANN to produce a source estimate based on simulated pairs of source and M/EEG activity and achieve high accuracy compared to many conventional methods (Cui et al., 2019; Hecker, Rupprecht, van Elst, & Kornmeier, 2020, 2022; Pantazis & Adler, 2021).

Multiple Signal Classification (MUSIC, Mosher & Leahy, 1998) and recursively applied (RAP-) MUSIC (Mosher & Leahy, 1999) are popular approaches for solving the M/EEG inverse problem. Both methods are based on the concept of subspace estimation, which involves estimating the subspace of the neural activity from the measured M/EEG signals using singular value decomposition (SVD). While MUSIC calculates an inverse solution by applying a SVD on the data covariance matrix in a single step, RAP-MUSIC extends the MUSIC method by repeatedly applying the MUSIC algorithm to the measured M/EEG signals. At each iteration, the estimated signal subspace is used to update the estimate of the neural activity, and the updated estimate is then used to update the estimate of the noise subspace. This recursive process continues until convergence is achieved. An improvement of the RAP-MUSIC algorithm was proposed by truncating the recursively calculated subspace with each iteration (TRAP-MUSIC, Mäkelä, Stenroos, Sarvas, & Ilmoniemi, 2018). This effectively removes residual variance that could not be explained in the prior iterations which leads to disturbances in localization (“RAP dilemma”, cf. Fig. 2 in Mäkelä et al., 2018).

Algorithms that follow the RAP-MUSIC-scheme have in common that a spatially coherent *source patch* will not be detected reliably. Approaches to find multiple dipoles per recursion were proposed by Mosher and Leahy (1998) and Katyal and Schimpf (2004), albeit with factorial increase in computational complexity. Liu and Schimpf (2006) introduced a more computationally efficient way to find extended source clusters by applying computing a weighted Minimum Norm Estimate (wMNE) solution on all estimated single dipoles and their respective neighbors. This step, however, can introduce (1) spurious sources at locations in which a single dipole would have explained sufficient parts of the signal subspace and (2) the approach is limited to finding sources of larger extent beyond single dipoles and their first-order neighbors.

In this paper we summarize the results of our endeavor to overcome this limitation in creating an analysis algorithm that is capable to effectively solve the inverse problem in constellations where established methods do have limitations.

## 3 Methods

The M/EEG inverse problem refers to the process of inferring the underlying neural activity that generates a given set of EEG measurements. This problem is a fundamental challenge in the field of neuroscience, as it involves understanding the spatio-temporal patterns of neural activity that underlie various cognitive and behavioral processes.

The EEG inverse problem is described as correctly identifying the source matrix *J* ∈ ℝ^*p×t*^ that, when multiplied by the leadfield matrix *L* ∈ ℝ^*q×p*^ (also referred to as gain-matrix) produces the observed EEG data matrix *M* ∈ ℝ^*q×t*^ with *q* electrodes, *p* dipoles (i.e., positions in the brain) and *t* time points.

The propagation of neural currents *J* through the leadfield *L* is thus defined as:

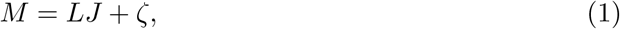

where *ζ* is the noise in the EEG data. For simplicity, we assume dipoles with fixed orientation perpendicular to the cortical surface.

### 3.1 Multi-Signal Classification (MUSIC) and its iterations

MUSIC approaches can be interpreted as algorithms that aim to select candidate dipoles in the brain that explain the signal of the EEG data. This is accomplished by calculating the signal and noise subspace of the data covariance *C* ∈ ℝ^*q×q*^ :

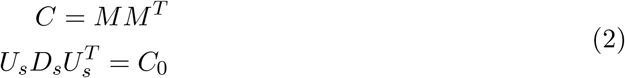

where *M* ^*T*^ denote the transpose of the EEG data matrix *M* and *C*_0_ denotes the covariance matrix of the noiseless data. *U*_*s*_ denotes the eigenvectors and *D*_*s*_ the eigenvalues that both belong to the signal subspace. Signal and noise are sought to be disentangled by selecting only the first *n* eigenvalues of the covariance matrix *UDU* ^*T*^ = *C*. While this selection is inherently difficult, it was recommended to overestimate *n* to avoid losing parts of the signal subspace (Mosher & Leahy, 1999). We followed a different approach by algorithmically selecting the set of eigenvalues belonging to the signal subspace. First, eigenvalues were normalized to a maximum of 1 by dividing all eigenvalues by the largest eigenvalue, yielding the normalized set of eigenvalues 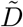. We then calculate the difference from each eigenvalue to the next eigenvalue, yielding 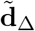. Let 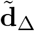 be the set of eigenvalues and *ϵ* = 0.01 be the relative selection criterion, which was determined empirically during our testing phase. The smallest eigenvector *ñ* that belongs to the signal subspace is defined as

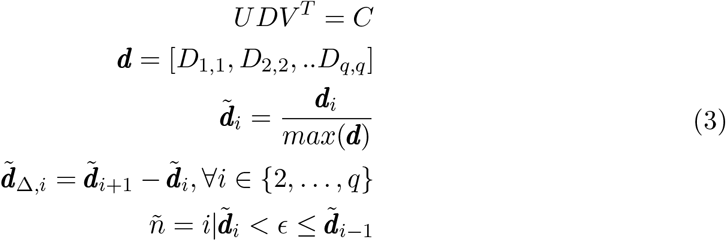

The estimated signal subspace is thus *U*_*s*_ = *U* (1 : *ñ*) and the projection to the signal space is defined as 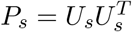.

The MUSIC localizer is then calculated as follows:

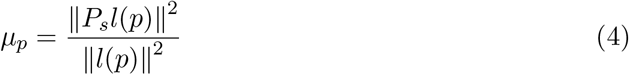

where l(p) denotes the *p*^*th*^ column of the leadfield matrix *L*. The resulting localizer *μ* is finally filtered to only contain values above a certain criterion (typically between 0.9 to 0.99).

Recursively applied MUSIC (RAP-MUSIC) also makes use of the signal subspace and its projection by iteratively selecting candidate dipoles as follows. We henceforth describe the RAP-MUSIC algorithm.

The first candidate is the dipole with the largest source amplitude in the MUSIC localizer as described above (eq. 4). Let *Î*_1_ be the topography of the initially selected candidate at iteration *i* we construct a set of topographies *B* = [*Î*_1_, …, *Î*_*i*_] that stores all topographies of the selected candidates. Using the set of topographies *B* ∈ ℝ^*q×i*^ at iteration *i* we define the out-projector matrix *Q*_*i*_:

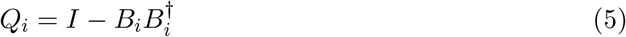

where *I* ∈ ℝ^*q×q*^ denotes the identity matrix and *B*^†^ denotes the Moore-Penrose pseudo inverse of *B*.

The updated covariance matrix *C*_*i*_ is then calculated by multiplying the out-projector matrix by the signal subspace and the new signal subspace is thusly calculated:

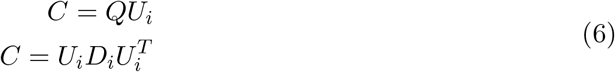

We then calculate the new signal subspace projection as *P*_*i*_ = *U*_*i*_(1 : *ñ*)*U*_*i*_(1 : *ñ*)^*T*^. Note, that the number of selected components can be truncated to 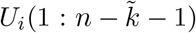 to alleviate the problem of the RAP-dilemma (Mäkelä et al., 2018). The (T)RAP-MUSIC localizer is then calculated as follows:

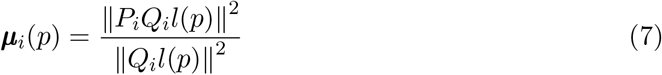

where l(p) denotes the *p*^*th*^ column of the leadfield matrix *L*. The dipole at which *u*_*i*_ is maximal is selected as the new candidate *c*_*i*_. The equations 5, 6 & 7 are iterated while *i < q* or a stopping criterion is met. The stopping criterion typically is met when the maximum of the (T)RAP-MUSIC localizer *u*_*i*_ falls below a threshold (in our case 0.975), i.e., when no dipole is able to explain the signal subspace projection sufficiently.

### 3.2 FLEX-MUSIC

A new dipole is selected in each iteration of the RAP-MUSIC algorithm based on the current subspace projection *P*. As described by Mäkelä et al. (2018), the selection of the most optimal dipoles often leaves some residual of the signal subspace projection to be explained. One reason for this is that neural sources can often not be represented by a single dipole due to the functional coherence in the cortex that is reflected by locally smooth activity of varying extent. To overcome this limitation we extended the set of candidate dipoles by multiple sets of smoothly distributed dipole clusters.

We have therefore calculated gradients *G*_*s*_ ∈ ℝ^*p×p*^ for increasing smoothness orders *s* ∈ 1, 2, …, *S*. Each gradient *G*_*s*_ transforms the original leadfield matrix *L* to leadfield matrices of increasing smoothness orders *L*_*s*_ as based on the neighborhood matrix *A* ∈ ℝ^*q×q*^ :

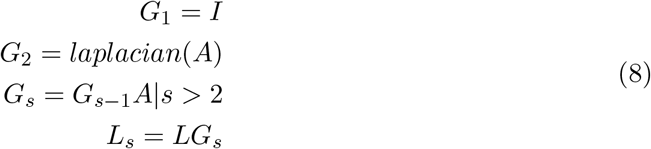

where S is the largest order of smoothness and *I* ∈ R^*p×p*^ is the identity matrix.

The selection of S depends on the upper boundary of smoothness to be assumed in the source model and should be adjusted depending on the number of dipoles in the source model. Note, that the original leadfield, denoted as *L*_1_, remains, since the respective gradient *G*_1_ is the identity matrix.

We calculate the FLEX-MUSIC localizer at iteration i by

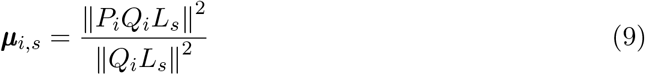

yielding one localizer for each smoothness order *s*.

The dipole or dipole cluster 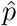 (depending on the estimated optimal smoothness order *ŝ*) at which ***μ***_*i,s*_ is maximal is selected as the new candidate.

We then update the set of topographies *B* with the newly added topography 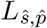. Furthermore, we update the source covariance matrix *S* by adding the column vector of 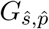:

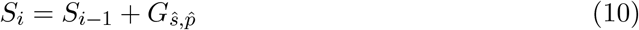

FLEX-MUSIC iterates the equations 5, 6, 8, 9 and 10. No truncation, as done in TRAP-MUSIC, was applied since that yields better results during testing.

Unlike MUSIC, the recursive approaches (e.g., RAP-, TRAP- and FLEX-MUSIC) require a final estimation of the current source density 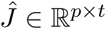 after candidate selection. We use the source covariance matrix *S* to calculate a weighted minimum-norm-like solution:

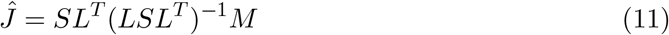

where *M* ∈ ℝ^*q×t*^ is the EEG or MEG data matrix.

In summary, FLEX-MUSIC further alleviates the RAP-dilemma of residual variance in the signal subspace projector *P*. This is achieved by adding clusters of neighboring dipoles to the set of candidate dipoles at each recursion step. The algorithm is still capable to localize single dipoles since they remain part of the set of candidates (cf. eq. 8). This renders FLEX-MUSIC a flexible solution to the M/EEG inverse problem in which the spatial extent of neural activations is often unknown *a priori*. In this way, FLEX-MUSIC increases the probability to identify extended sources and integrate their extensions into the result source space, instead of either ignoring the extension or treating the neighboring dipoles as separate sources.

### 3.3 Evaluation

In order to evaluate the proposed method, we simulated pairs of source- and EEG-data using an anatomical template brain “fsaverage” (Fischl, Sereno, Tootell, & Dale, 1999) by the Freesurfer image analysis suite^1^. EEG simulations were carried out using a precomputed three shell boundary element method (BEM; Fuchs, Kastner, Wagner, Hawes, & Ebersole, 2002) forward solution as provided by mne-python (v20.3, Gramfort et al., 2013). Each shell (brain, skull & scalp tissue) was composed of 5120 vertices. The conductivity was set to 0.3*S/m*^2^ for brain and scalp tissue, and 0.06*S/m*^2^ for the skull.

The source model was chosen with *p* = 1, 284 dipoles with icosahedral spacing. For the EEG electrodes we used the Biosemi 64-channel layout consisting of *q* = 64 electrodes of the 10-20 system. Using the forward model and the parameters described, we calculated a leadfield *L* ∈ ℝ^*q×p*^.

We evaluate our proposed method FLEX-MUSIC by comparing it to a diverse set of other inverse algorithms including TRAP-MUSIC, eLORETA, a sparse Bayesian learning (SBL) approach called Convexity Champagne and the Multiple Constrained Minimum Variance Beamformer (MCMV, Cai et al., 2022; Mäkelä et al., 2018; Mosher & Leahy, 1999; Nunes et al., 2020; Pascual-Marqui, 2007; D. Wipf & Nagarajan, 2009).

Motivation is given for the choice of each method for solving the EEG inverse problem. TRAP-MUSIC was chosen as one of the latest developments of the recursive MUSIC approaches. eLORETA is a popular choice in many EEG studies with theoretically low localization errors, rendering it the most suitable choice within the minimum-norm family. Convexity Champagne was chosen as a very recent improvement to the Champagne algorithm. Champagne was shown to produce fast and accurate solutions within the framework of *empirical Bayes* (D. P. Wipf et al., 2010). MCMV is a similar approach as the linearly constrained minimum variance (LCMV) beamformer. However, it is designed to be less prone to correlations between sources, rendering it a useful innovation over LCMV. All methods were implemented in-house and are available in our python package *invertmeeg*^2^.

Optimal regularization of eLORETA, MCMV and Convexity Champagne was achieved using generalized cross validation (GCV, Grech et al., 2008) on a set of 7 regularization parameters *λ* = {10^−3^…, 10^3^}. The MUSIC-type methods (FLEX- and TRAP-MUSIC) do not require the regularization parameters since the noise is estimated by selecting the signal subspace.

A set of 1,000 samples consisting of ground truth sources *J* ∈ ℝ^*p×t*^ and corresponding EEG *M* ∈ ℝ^*p×t*^ was simulated in order to evaluate the accuracy of all solvers.

The number of simulated consecutive time points was set to *t* = 20. The simulation parameters are outlined in Tab. 1. Half of all samples contained single dipoles, whereby the other half contained samples of extended dipole clusters with coherent activity over time. The cluster size was varied in terms of neighborhood orders, whereas an order of 1 indicates a single dipole and an order of 2 indicates a dipole including all its neighbors. The source time course was generated as random sequence of a colored frequency spectrum as described by the 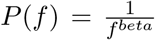, where beta controls the level of temporal smoothness. Noise was generated as random spatio-temporal white noise with inter-channel correlation between -1 and 1. The noise was added to the EEG matrix such that a random signal-tonoise ratio (SNR) within the given range outlined in Tab. 1 was achieved. Only white noise is considered since the presence of colored noise in real data can be handled by whitening the EEG data as a preprocessing step.

**Table 1:**
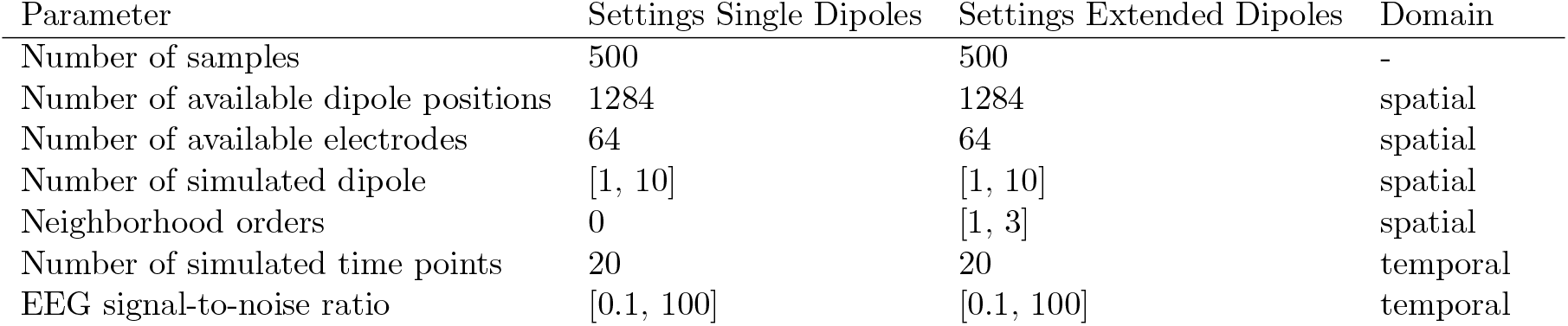
Simulation Parameters. Parameters and parameter ranges (denoted in square brackets) for the source and EEG simulations of single and extended dipoles. Diameters are reported in neighborhood orders (the higher, the larger the diameter of the cluster).

We calculated inverse solutions to each of the simulated samples of EEG data using the different methods, as described above. Accuracy of the individual inverse solutions is quantified by calculating the mean localization error (MLE), Earth Mover’s Distance (EMD, Hitchcock, 1941) and the mean squared error (MSE). Furthermore, we quantified the sparsity of each inverse solution.

We calculated the MLE by first identifying the local maxima of the ground truth source matrix *J* and the inverse solution 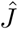 and then calculating the minimum Euclidean distance between each true dipole location and all estimated dipole locations.

MSE quantifies how close the estimated dipole moments (in *nAm*) are to the true dipole moments and was calculated as follows:

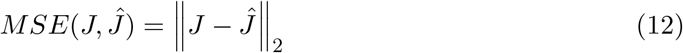

EMD is a measure that calculates the distance between two distributions. It is a suitable method to quantify the accuracy of an inverse solution since, unlike MSE, it takes into account the distance between dipole locations. It was calculated as follows:

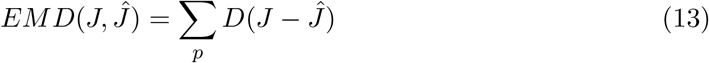

where *D* ∈ ℝ^*p×p*^ is the distance matrix containing the Euclidean distance between each dipole pair. Prior to calculating the EMD, we have computed the absolute mean over time points for *J* and 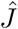 and normalized them.

Sparsity was calculated by first normalizing the columns of the estimated source matrix 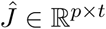 to unit length by division of the respective columns L2-norm. The L1 norm of the normalized matrix 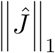 was then calculated, yielding a metric with an inverse relationship to sparsity. According to the dogma of Occam’s Razor, which states that complexity should not be posited without necessity, we can assume that sparse solutions make the fewest assumptions about the brain’s activity and are therefore preferred, given that they explain a sufficient amount of the data.

In summary, the evaluation metrics described above capture the accuracy of estimated local maxima positions (MLE), the accuracy of the global pattern of the inverse solution (EMD), the accuracy of dipole moments (MSE) and the sparsity of the solution (L1 norm).

## 4 Results

We calculated the accuracy of all inverse algorithms as described in the previous section. Exemplary samples of ground truth source activity and estimated sources are shown in Figure 1.

**Figure 1:**
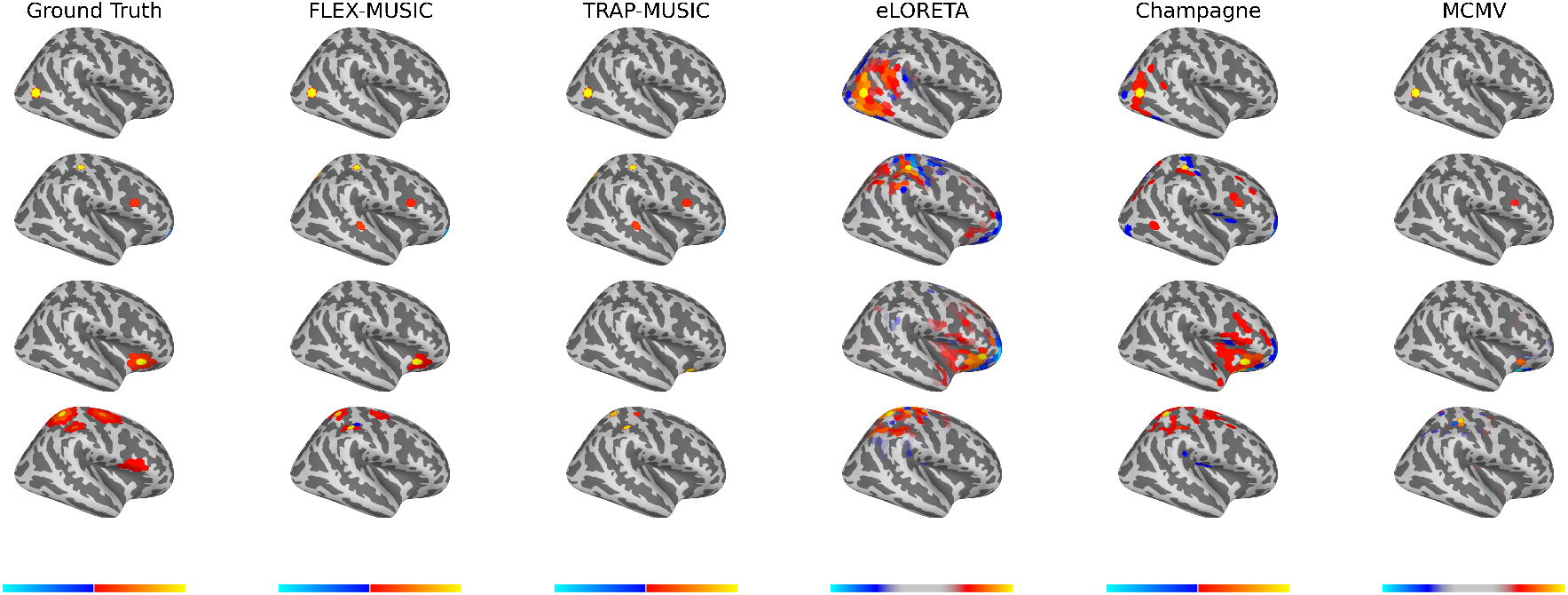
Examples of ground truth and estimated sources. Exemplary plots of ground truth sources and the estimated sources of FLEX-MUSIC and comparative approaches. For demonstration purposes, samples were selected based on visibility on the right lateral view. First row: Single non-extended source. Seconds row: Multiple non-extended sources. Third row: Single extended source. Fourth row: Multiple extended sources. Colorbars were adjusted for eLORETA and MCMV to improve visibility of the source pattern. FLEX-MUSIC visibly recovers the actual source extent whereas all other approaches tested show biases towards single dipoles or extended dipoles.

Figure 2 depicts the Mean Localization Error (MLE), Earth Mover’s Distance (EMD) and the Mean Squared Error (MSE) for all inverse solutions of each solver. Samples were divided into those containing single dipole sources and those containing extended source clusters. FLEX- and TRAP-MUSIC show overall lowest MLE and EMD for single dipole sources when compared to all other solvers. Notably, the median MLE is zero for both MUSIC-based methods, and there was no significant difference in MLE for single-dipole sources (*p* = 0.96, *t* = 0.36, *d* = 0.02). FLEX- and TRAP-MUSIC did also not differ significantly in EMD (*p* = 0.95, *t* = 0.73, *d* = 0.05) and MSE (*p* = 1.00, *t* = 0.31, *d* = 0.02). Champagne produced the next best accuracies, eLORETA and the MCMV Beamformer show comparatively poor accuracy in correctly estimating the source distribution as depicted in relatively high EMD and MSE, whereas the Convexity Champagne solver lies in-between.

**Figure 2:**
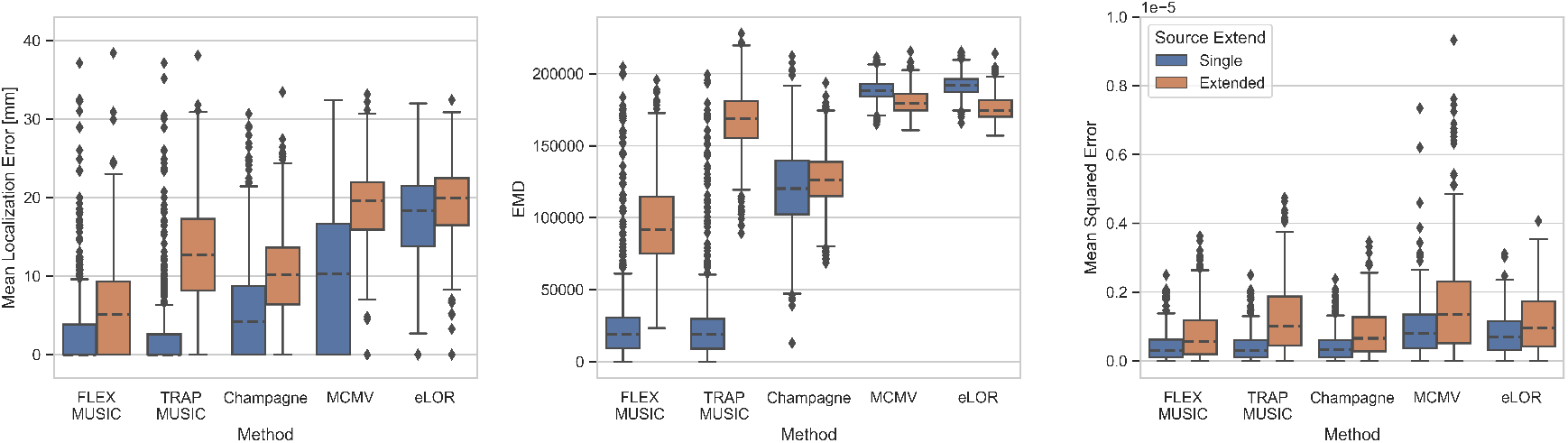
Evaluation of all solvers. Boxplots depict the accuracy of FLEX-MUSIC and all other solvers tested. Left: Mean Localization Error in mm. Center: Earth Mover’s Distance (EMD), Right: Mean Squared Error. Blue: Single dipoles. Orange: Extended dipoles. Note, that FLEX-MUSIC achieves competitive accuracy for single-dipole sources and the highest accuracy for samples containing spatially extended sources.

The advantage of the proposed FLEX-MUSIC solver becomes most apparent for extended sources. While TRAP-MUSIC fails to accurately localize sources with spatial extent, FLEX-MUSIC retains the lowest MLE and EMD compared to all other solvers (Fig. 2).

Figure 3 shows the sparsity of the produced inverse solutions of all solvers, defined as the L1-norm of the L2-normalized source estimate 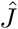. We find that FLEX- and TRAP-MUSIC exhibit the highest sparsity, MCMV and eLORETA the lowest, and Convexity Champagne lies in the middle. Interestingly, only FLEX-MUSIC shows a clear difference in sparsity between samples containing single dipoles and those containing extended dipole clusters, which is consequence of its flexibility to estimate sources of varying extent.

**Figure 3:**
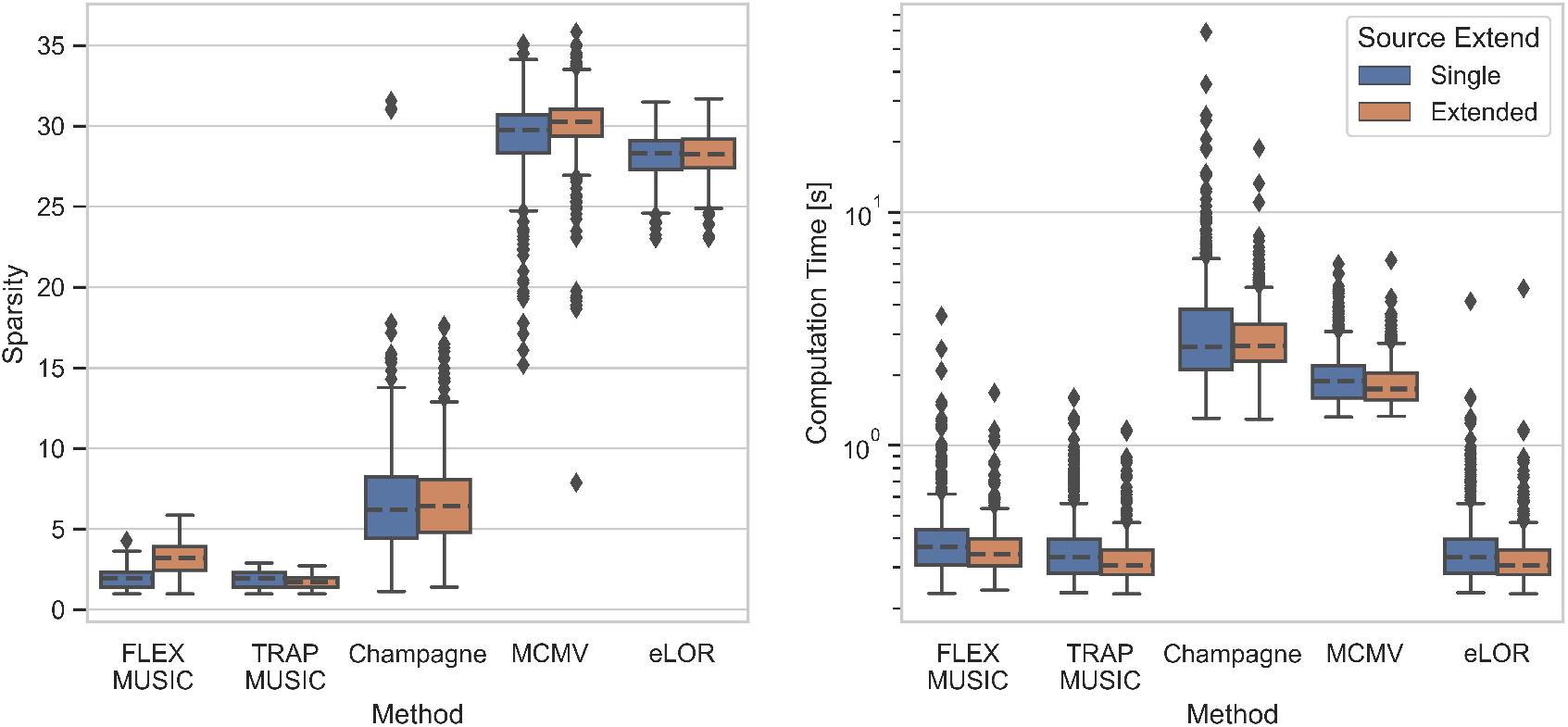
Sparsity of inverse solutions and time of computation. Boxplots depict the L1 norm (left) and the computation time in seconds (right) for all inverse solvers tested. The L1 norm reflects the level of “non-sparsity”. Convexity Champagne requires ten times longer computation time compared to FLEX-MUSIC.

This aspect is shown in detail in Figure 4, depicting each solver’s capability to recover the spatial level of sparsity in the ground truth. FLEX-MUSIC shows the highest correlation (*r* = 0.82) between the sparsity in the ground truth sources and the sparsity in the predicted sources. TRAP-MUSIC shows a high correlation for highly sparse samples and is biased for less sparse samples. Convexity Champagne produced solutions that were often less sparse that the ground truth, whereas MCMV and eLORETA had a strong bias towards finding less sparse activations. Despite the strong bias, the sparsity of eLORETA was significantly correlated with the sparsity in the ground truth samples.

**Figure 4:**
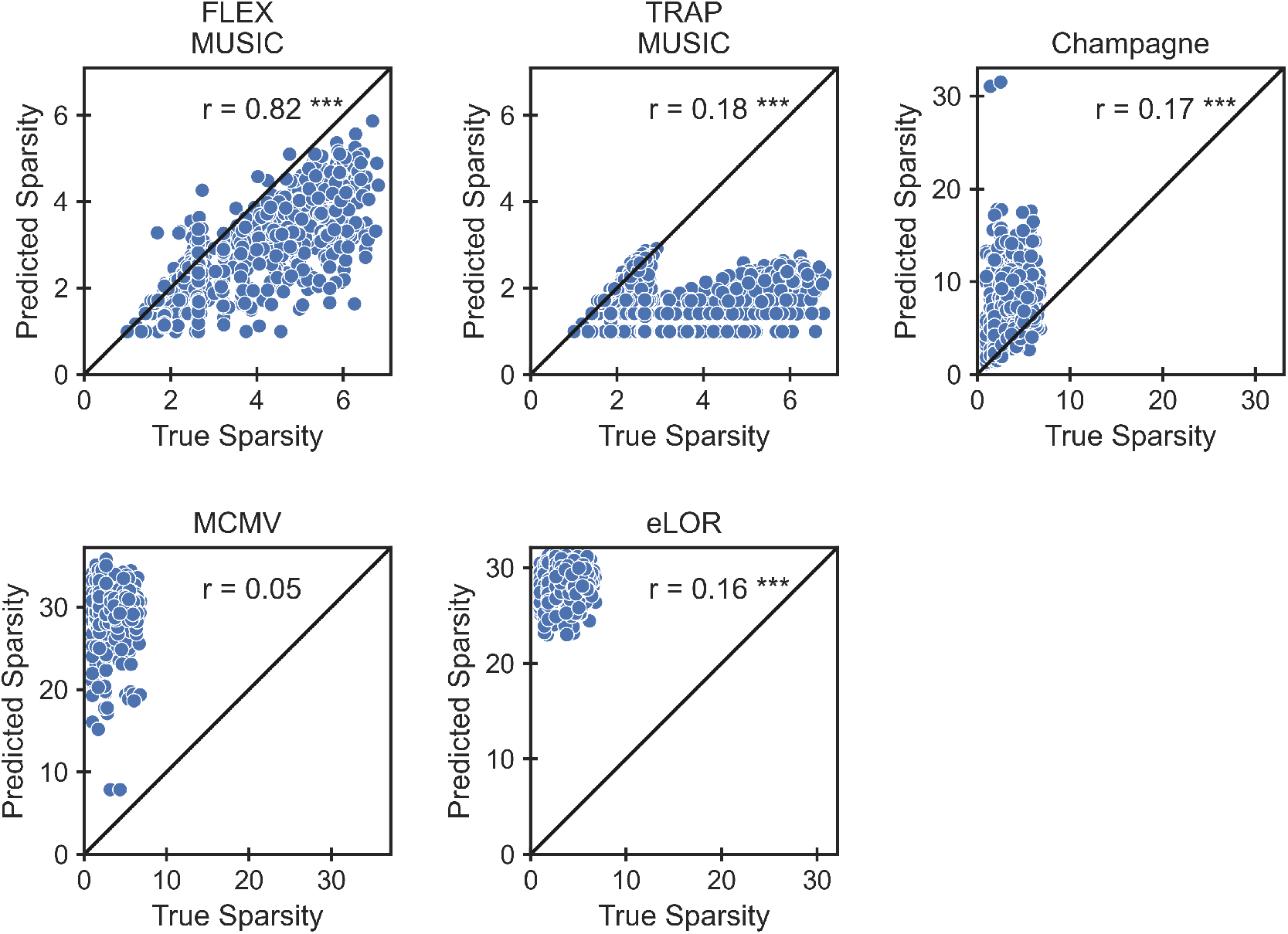
Accuracy in extent estimation. Scatter plots depict the relationship between the L1 norm in the ground truth and the L1 norm in the prediction. The L1 norm reflects the level of “non-sparsity”. Note, that only FLEX-MUSIC is capable to reproduce the true sparsity with high correlation (*r* = 0.82). \**p <* 0.05, \*\**p <* 0.01, \*\*\**p <* 0.001

Next, we tested the dependence of the inverse solution accuracy on varying levels of noise. For this comparison we have combined samples containing single dipoles and those containing extended dipole clusters (Fig. 5). The graph shows that FLEX-MUSIC yields inverses solutions with the highest accuracy regardless of the level of noise in the EEG data.

**Figure 5:**
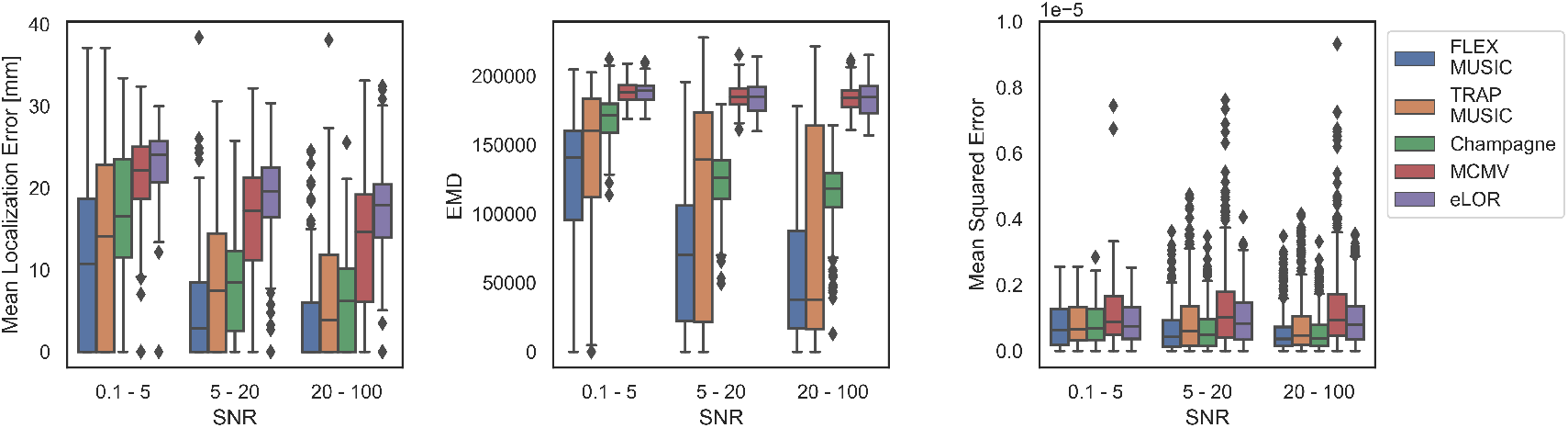
Dependence on SNR. Accuracy of all solvers separated by the signal-to-noise ratio (SNR) in the simulated EEG samples.

Finally, we analyzed the accuracy of all solvers depending on a varying number of dipoles/ dipole clusters in the ground truth. FLEX-MUSIC achieves again the highest accuracy on all metrics regardless of how many dipoles/ dipole clusters were present in the ground truth source.

### 4.1 Computational Expense

The computational expense to calculate the EEG inverse operators is presented in Figure 3 (right). Although FLEX-MUSIC has some additional processing steps compared to TRAP-MUSIC, we find that the median computation time differ only slightly (Δ*t* = 0.042*s, p* = 1.72·10^−6^, *t* = 5.25, *d* = 0.23). eLORETA required similar computation times of 0.32*s*. Convexity Champagne required the longest computation time of 2.66*s*. Note, that eLORETA, Convexity Champagne and MCMV were re-computed 7 times for varying levels of regularization. If the optimal regularization parameter was known in advance, the computation time could be reduced seven-fold.

**Figure 6:**
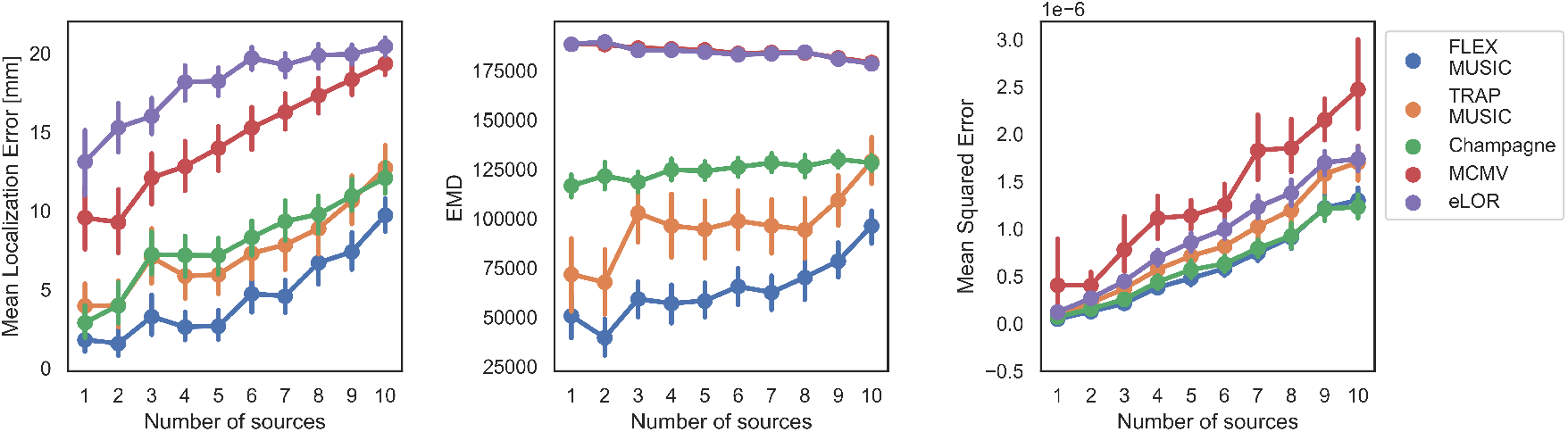
Dependence on number of active sources. Accuracy of all solvers for varying numbers of active sources within the simulated EEG samples. Error bars depict standard errors of the mean (SEM).

**Table 2:** Medians of all metrics. Table depicts the median performance of each solver in each metric considered. Both single-dipole and extended-dipole samples were included in this analysis. Best performance per metric is highlighted in bold font. MLE: Mean localization error. EMD: Earth Mover’s Distance. Sparsity: Sparsity of the produced inverse solutions. Time: Computation time of the inverse operator.

| Method | MLE [mm] | EMD | Mean Squared Error | Sparsity | Time [s] |
| --- | --- | --- | --- | --- | --- |
| FLEX-MUSIC | **2.11** | **66,291.31** | **4.15e-07** | 2.36 | 0.35 |
| TRAP-MUSIC | 5.50 | 125,233.79 | 5.37e-07 | **1.73** | **0.32** |
| Champagne | 7.73 | 123,775.42 | 4.67e-07 | 6.29 | 2.66 |
| eLOR | 19.05 | 185,186.10 | 8.16e-07 | 28.29 | **0.32** |
| MCMV | 16.25 | 184,793.86 | 9.65e-07 | 30.04 | 1.82 |

## 5 Discussion

In this work we have presented FLEX-MUSIC, a new approach to solve the M/EEG inverse problem embedded in the recursive MUSIC scheme. Similar to the well-known RAP-MUSIC techniques, it iteratively adds candidate positions to the set of active sources. In addition to the RAP-MUSIC approaches it adds an extended dictionary of increasingly smooth source patches.

We have shown that FLEX-MUSIC works as well as TRAP-MUSIC in scenarios where the EEG was produced by singular dipole sources. Furthermore, we have shown that FLEX-MUSIC accurately estimates the spatial extent of the underlying sources. Strikingly, of all inverse solvers tested, only FLEX-MUSIC was capable of estimating the level of sparsity in the ground truth, showing the highest correlation between the sparsity levels in the ground truth sources and the estimated sources or *r* = 0.82, whereas the next best solver achieved only a correlation of *r* = 0.18. We consider this aspect to be the most compelling argument for FLEX-MUSIC, since the correct estimation of the extent of neural generators underlying an M/EEG signal is of high interest for multiple reasons.

First, as stated above, RAP-MUSIC approaches suffer in general from what is called the *RAP dilemma*, i.e., the interference of residual variance from previous iterations. Since FLEX-MUSIC is capable to explain a larger portion of the subspace, it effectively diminishes the residual variance from the previous iterations, leading to more accurate inverse solutions throughout all iterations.

Second, good estimations of the extent of neural sources is of high interest in the presurgical diagnostic in epilepsy. Not only does FLEX-MUSIC more reliably find the true location of the source maxima regardless of their extent (cf. Fig. 2), it is also capable to isolate the prospective resection area in the brain.

As stated earlier, the recursive MUSIC approaches are a suitable option to solve inverse problems where the solution space is considerably large due to their low computation time and built-in regularization. While the presented Convexity Champagne has shown competitive accuracy in many cases, we showed that the computational expense was almost 10 times higher in our setting. We further argue that the difference in computational expense may further increase with larger source models (i.e., higher number of dipoles), rendering the computation of Bayesian inverse solutions unfeasible fir certain clinical applications.

A potential weakness of our proposed FLEX-MUSIC algorithm is that mesoscale brain activity may not be sufficiently modelled with single dipoles and smooth dipole clusters. Various shapes, e.g., elliptical coherent sources, may be involved in real-world M/EEG recordings of brain activity. However, we expect that for a sufficiently large source model, FLEX-MUSIC should still be able to reconstruct deviant shapes of sources using the circular smooth patches. Future improvements of the FLEX-MUSIC algorithm could involve changing the way the dictionary of candidate dipoles or dipole clusters is applied. One potentially fruitful modification may be to find the set of active dipoles that explain the signal subspace in each iteration is found in a data-driven manner, unlike the dictionary-like search that we proposed in the present work. Another idea would be a combination of SBL and FLEX-MUSIC, in which a small set of active dipoles are added to the source covariance matrix (cf. eq. 10) based on the maximum a posteriori (MAP) estimation. An interesting development of the recursive MUSIC algorithms was presented recently by Adler, Wax, and Pantazis (2022), called Alternating Projections (AP). The approach yielded lower MLE compared to RAP- and TRAP-MUSIC, especially under low SNR conditions. It may be promising to translate the idea of FLEX-MUSIC to the domain of AP, which could potentially further increase the accuracy of the AP-approach in scenarios where spatially coherent sources are expected.

A very important finding of the present work, beyond the performance benefits of FLEX-MUSIC, is that different inverse solutions applied to the same data set can produce quite divergent results. In the case of real EEG data, where a ground truth is not available, we strongly recommend to use different methods for EEG source calculations in parallel and compare the results with each other. As we have shown in this work, combining FLEX-MUSIC and a Champagne algorithm may be useful, although Champagne may overestimate the size of neural sources in some cases. An interesting family of recently developed inverse solution are based on artificial neural networks (ANNs, Hecker et al., 2020, 2022). In a next step we plan to compare a potentially further optimized version of FLEX-MUSIC with this family of ANN-based solvers.

## Footnotes

1 http://surfer.nmr.mgh.harvard.edu/

2 https://github.com/lukethehecker/invert

